# A new microfluidic device for simultaneous detection of enzyme secretion and elongation of a single hypha

**DOI:** 10.1101/2022.04.07.487578

**Authors:** Ayaka Itani, Yosuke Shida, Wataru Ogasawara

## Abstract

Fungal cells perform enzyme secretion and elongation by exocytosis in the apical region. The widespread branching of hyphae and the inability to control environmental conditions over long periods make it difficult to observe and analyze single hyphae with conventional assays. Therefore, although hyphal morphology is closely linked to productivity, no real-time measurements of morphology and exoenzymes have been carried out. In this study, a microfluidic system was developed to compartmentalize a single hypha germinated from a single spore. This allowed detailed observation of a single hypha and provided new insights, such as the fact that enlarged vacuoles inhibit nuclear movement. Furthermore, a cellulase detection assay based on subtle differences in molecular polarity was developed to detect hyphal growth and enzyme secretion in real-time using *Trichoderma reesei*, a potent cellulase-producing hypha, as a model. When the fluorescence from the detection assay was compared with the GFP fluorescence intensity using a strain fused with cellulase CBHI and GFP, a strong correlation was observed. As *T. reesei* secretes a series of cellulases, these results prove that the extracellular enzymes can be measured in real time. This microfluidic system has enabled real-time visualization and analysis of cellular heterogeneity, hyphal and enzyme dynamics associated with carbon source exchange, and quantitative dynamics of gene expression. The technology can be applied to a wide range of other biosystems exhibiting similar polar cell growth, from bioenergy production to human health.

**IMPORTANCE:** Hyphal morphology and productivity of filamentous fungi are linked by exocytosis. Conventional assay methods make it difficult to observe and analyze single hyphae. Here, a robust and high-performance microfluidic system was developed to compartmentalize single hyphae germinated from a single spore, enabling their long-term observation. Using the potent cellulase-producing fungus *Trichoderma reesei*, the system made it possible to visualize and analyze cell heterogeneity, hyphae, enzyme dynamics, and quantitative gene expression dynamics associated with carbon source exchange. The technique can be immediately extended to various other biosystems exhibiting similar polar cell growth and is expected to contribute significantly to the elucidation of filamentous fungi production systems.

## INTRODUCTION

Filamentous fungi are used as microbial cell factories to produce antibiotics, organic acids, and enzymes (1). Many of these products constitute a multi-billion-dollar industry, the value of which is expected to increase with the transition from petroleum to a bio-based global economy (2). Filamentous fungi have attached growing interest for their utility in bioethanol fermentation as they can grow on a wide range of sugars and are highly resistant to many of the inhibitory molecules produced by hydrolyzed plant biomass (3, 4).

The filamentous fungus *Trichoderma reesei* (anamorph of *Hypocrea jecorina*) is a fast-growing conidiophore fungus widely distributed in soil environments. *T reesei* is known to secrete large amounts of cellulolytic enzymes (5) and has been extensively studied as a model fungus to understand the mechanism of cellulase production. The biosynthesis of cellulases in *T. reesei* is controlled by a complex network involving multiple regulatory proteins at the transcriptional level, and little is known about how cellulase genes are regulated and how the expression level of each cellulase gene is determined (6, 7). In Japan, mutants of *T. reesei* have been intensively studied after the country experienced a series of oil shocks in the 1970s (8). Their comparative genomic analyses have not only paved the way for the development of technologies for the industrial exploitation of *T. reesei* but have also shown that they hold the key to solving the regulatory puzzle of cellulase induction (9, 10) (Fig. 1).

**Figure 1.**
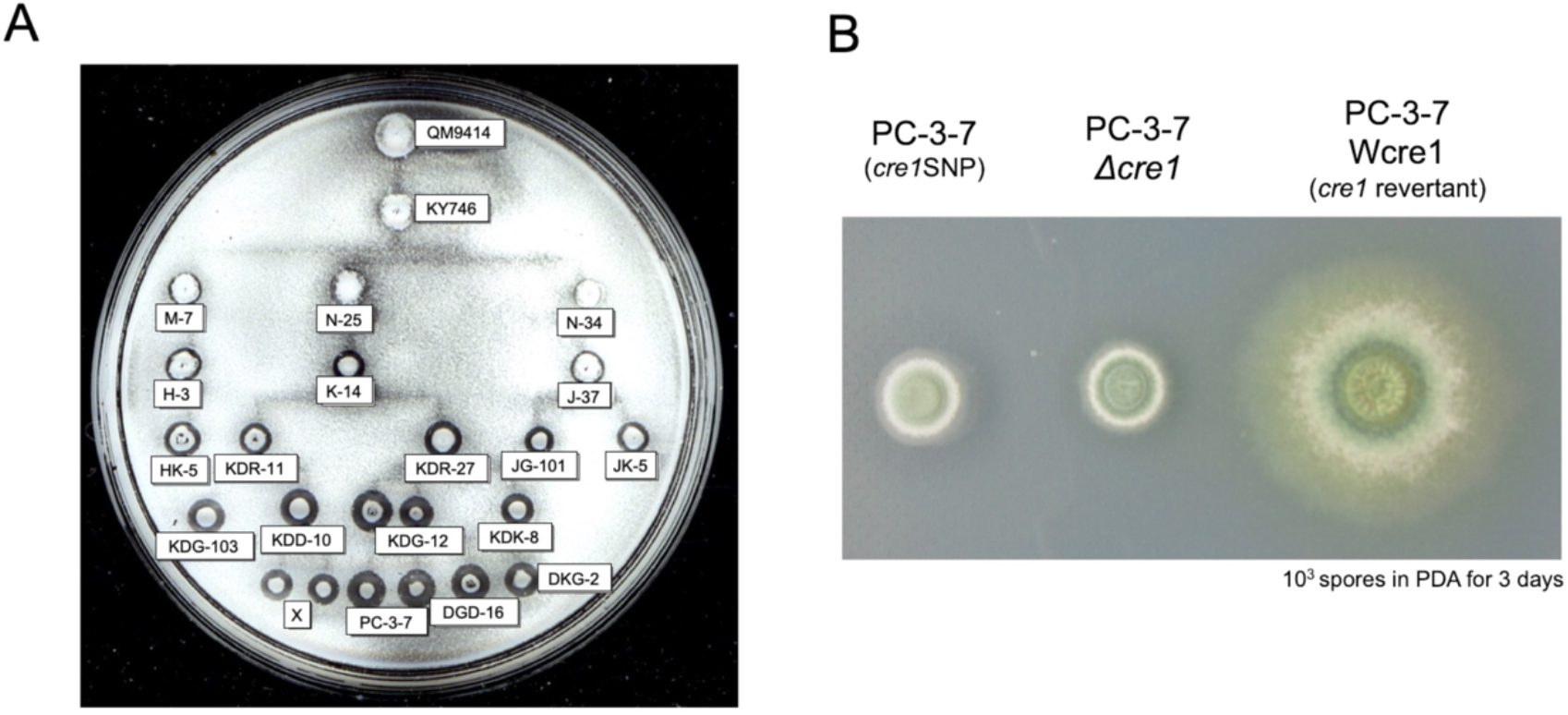
Mutant strains of *T. reesei* developed in Japan and morphological changes caused by CRE1, a gene involved in both morphology and cellulase production. (A) Conidia from each mutant were inoculated onto an agar plate containing cellulose as a carbon source. The clearing zone around the colony shows cellulose degradation (8). (B) PC-3-7 and PC-3-7 *cre1* revertant (PCWcre1) strains differ by a single nucleotide in the whole genome, but cause significant morphological changes, suggesting that CRE1 has other functions besides the regulation of cellulase expression.

The morphology of filamentous fungi plays an important role in determining their productivity. Specific morphology is needed for optimal productivity (11, 12), and the optimal morphology is different for each bioprocess (13). Therefore, strain engineering to optimize the morphological parameters for biotechnological applications has become a major goal of applied fungal research (1, 2). The shape of the hypha is determined by the assembly of the cell wall (14), which is a polar process that takes place at the tip. Increasing the number of hyphal tips is thought to be useful for improving protein secretion, and there is also some evidence supporting this hypothesis (15, 16). However, an increase in the number of hyphal tips may not correlate with an increase in protein titer (17-19). In other words, while it is important to investigate the relationship between micro/macromorphology and productivity, there is a growing demand to identify the growth and morphology of the hypha itself and its productivity.

Previous studies on single hyphae have mainly used methods in which hyphae are randomly placed on glass substrates or agar blocks and observed under a microscope (20-22). However, these methods do not provide temporal and spatial control over hypha tracking because the hyphae are extensively branched and cannot be cultured for long periods. To quantify the dynamics of large cell populations and to understand cell-to-cell variability, new approaches that can confine large numbers of hyphae along parallel pathways and that can control the microenvironment are needed.

Microfluidics is a powerful and excellent tool for manipulating and analyzing single cells, as it can confine cells within microstructures with dimensions comparable to those of cells (23-25). Here, we have developed a microfluidic platform for capturing single hyphae to grow them in spatially separated channels and for performing comparative analysis of hyphal growth and dynamics. In combination with live-cell imaging and fluorescent fusion protein technology, we investigated the growth of single hyphae, the organization and movement of nuclei, and the dynamics of cellulase gene expression over time in response to carbon source switching.

## RESULTS

### Microfluidic device design

We used multilayer soft lithography to fabricate a “conidia trap” microfluidic device (Fig. 2A and B) composed of polydimethylsiloxane (PDMS), which allows the observation of a single hypha in real time. The device can be divided into three main regions. The first region is the conidia loading channel (width 200 μm × height 30 μm, Fig. 2C and D, green) with a “conidia inlet” for the introduction of conidia and a “conidia outlet” to release the excess of conidia. The second area is a channel with a “bottle neck” design for conidia capture (conidia trapping site: length 7.0 μm × width 7.0 μm, bottle neck: length 5.0 μm × width 3.5 μm, Fig. 2D, light blue), allowing the observation of a single hypha extension into the observation channel (length 3.0 mm × a series of width). The widths of the channels were set from 5.0 μm to 17.5 μm with increments of 2.5 μm. The hypha observation channel (blue) was set to a height of 7.0 μm that was the maximum diameter of the hypha, to keep the hypha in focus. The third area is a channel (200 μm wide × 30 μm high, Fig. 2B Grey) with a “medium inlet” for the introduction of medium and a “medium outlet” to release the medium and waste.

**Figure 2.**
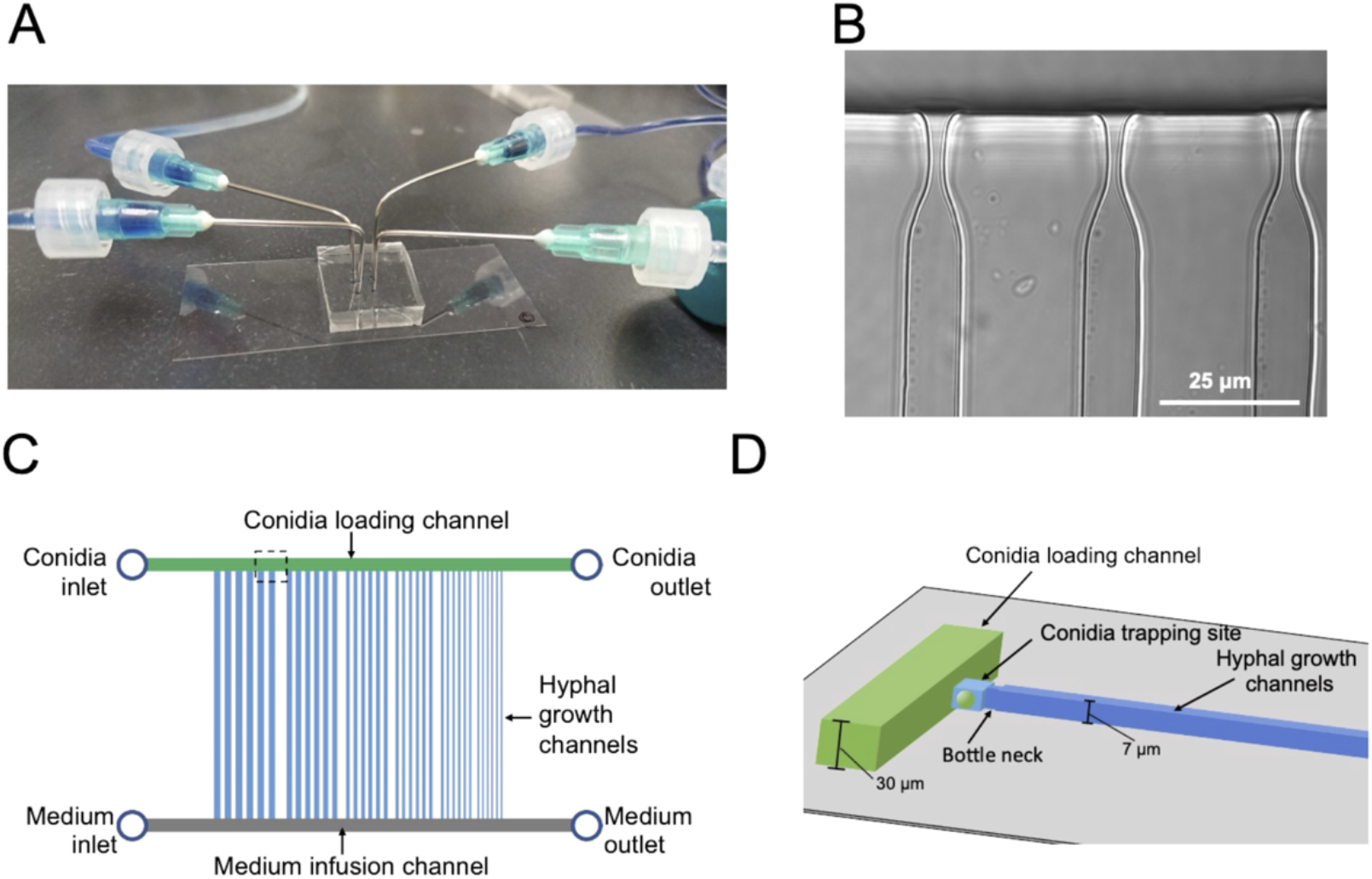
A microfluidic device for analyzing the growth and dynamics of a hypha. (A) External view of the microfluidic device. (B) Image of the constricted structure for conidia trapping. Scale bar: 25 μm. (C) Schematic representation of the chip layout: it consists of two inlets for supplying cells and medium, respectively, two outlets, a parallel array of conidia supply channels (green), medium injection channels (gray), and hypha growth channels (blue). (D) Schematic representation of the conidia trapping site containing two short channels of different dimensions (light blue) connecting the conidia supply channel (green) and the hypha growth channel (blue). The narrower and shallower channel traps a single conidium.

### Characterization of molecular diffusion in the microfluidic device

To ensure that the medium required for hyphal growth was diffused into the device, CFW was injected through the medium inlet at a flow rate of 0.5 mL/h after sufficient injection of carbon-free medium (basic medium). All channels diffused at approximately the same time (Fig. 3). The slightly faster rate in the narrower channels is due to the hydrodynamic properties: the closer the growth channel is to the medium inlet, the faster molecular diffusion starts, and the narrower the growth channel, the faster molecular diffusion is completed.

**Figure 3.**
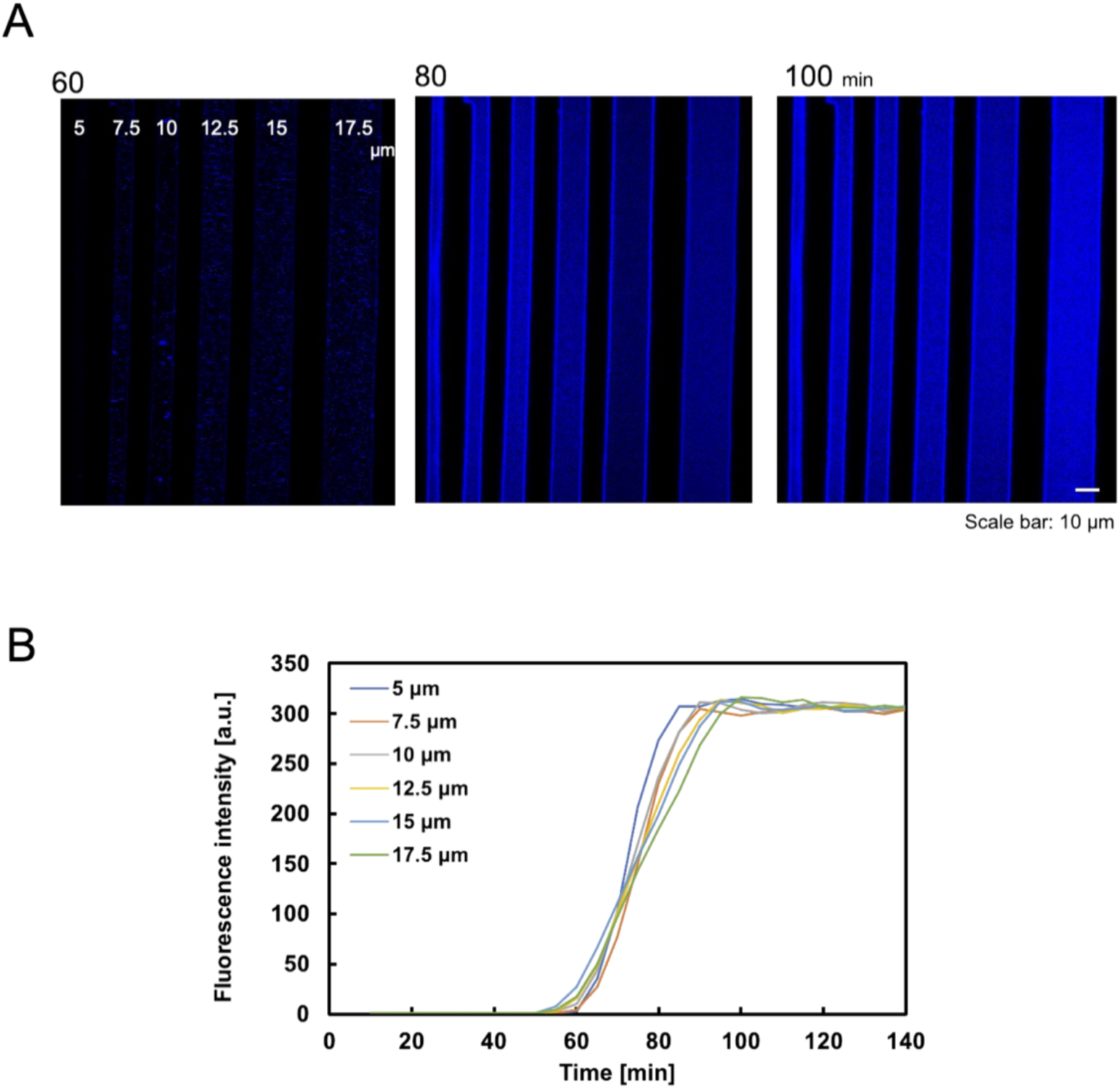
Molecular diffusion properties on microdevices with different channel widths. (A) Visualization of the gradient profile. Scale bar:10 μm. (B) Quantification of the gradient profile of CFW diffusion to steady state. (The measurements were carried out in the middle of the channel, just below the spore trap site.)

### Trapping of single conidia and hyphal growth

Since the apical tip of the hypha is the elongation point and the site of enzyme secretion, this device was designed to trap conidia at the entrance of the hyphal growth channel and to observe the hyphal elongation. The conidia of *T. reesei* have an initial diameter of about 3 μm and they swell to about 10 μm before germination. Therefore, the conidia were inoculated into the basic medium containing 1% glucose, left at room temperature for 5-6 hours to swell to about 5 μm, and washed in the basic medium without the carbon source. The swollen conidia were then suspended in the basic medium and introduced into the conidia inlet using a syringe. Conidia with delayed swelling and dead conidia slipped through the bottle neck structure and were ejected from the medium outlet. Consequently, most of the trapped conidia germinated. The germination time of conidia varied greatly, but the selective trapping made it possible to match the germination time to some extent. In addition, the wide constriction prevented the polarity anomaly of the filamentous fungus (Fig. 4A).

**Figure 4.**
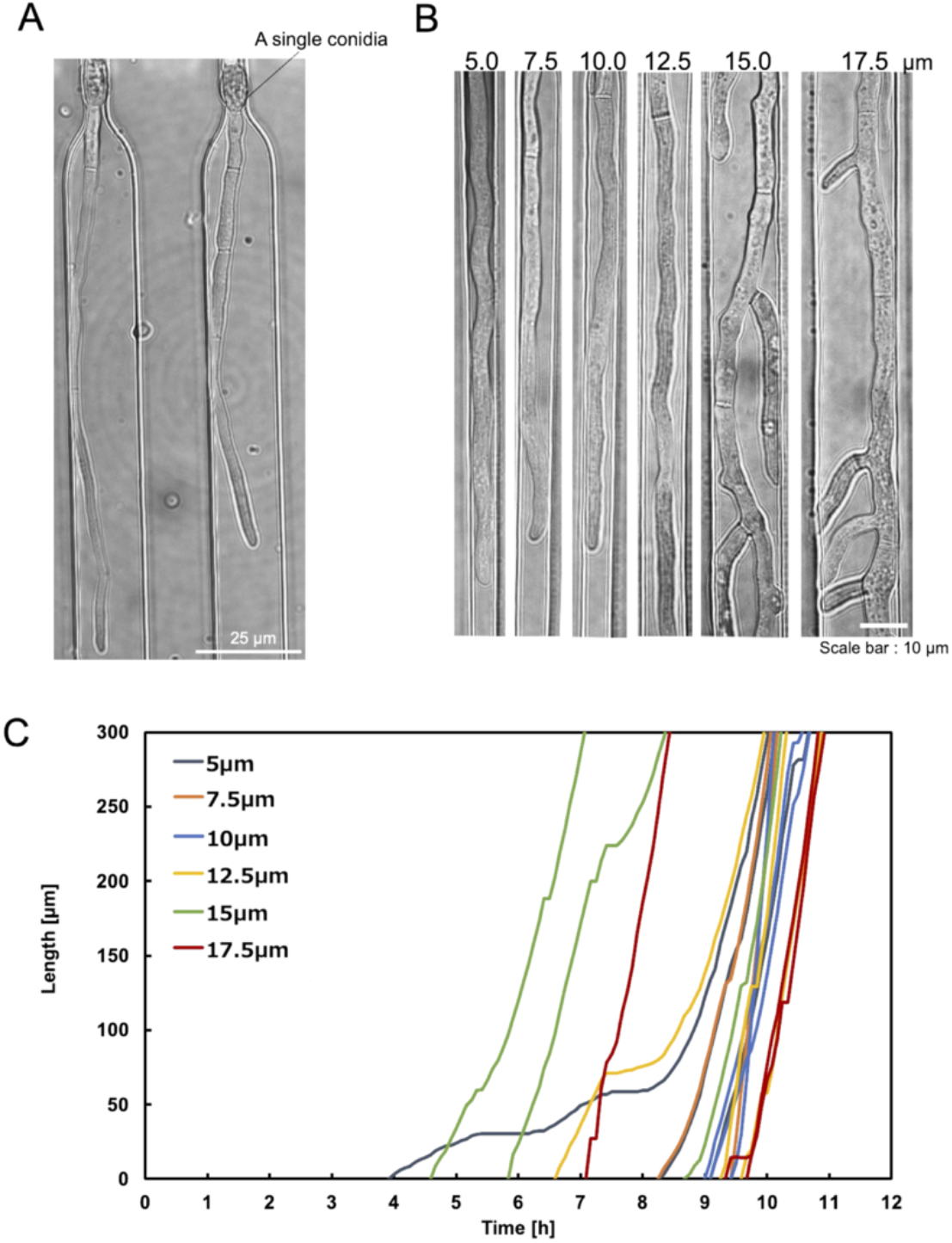
Single conidia trapping and compartmentalized hyphal growth. The hyphae are cultivated under a constant flow of 1% glucose in the basic medium at 0.5 mL/h. (A) A single conidium of *T. reesei* was trapped at the conidia trapping site. Scale bar: 25 μm. (B) The compartmentalized hyphal extension of *T. reesei* along the hyphal growth channels. Observation at 11 hours. Scale bar: 10 μm. (C) Time course of three individual hyphal length elongation per channel width on a single device. The hyphal length is measured at regular intervals of 5 min starting after conidia trapping.

After the conidia loading channel had been filled with conidia suspension, the conidia outlet was closed and pressurized with a syringe. In this way, a single conidium was retained in the conidia trap structure (Fig. 4A). Once the trapping site was occupied, subsequent conidia could preferentially travel to the next empty trap structure. Using this method, a trapping efficiency of over 70% could be achieved. To induce the direction of conidial germination and hyphal growth into the medium infusion channel of the medium, the conidia inlet and conidia outlet were closed after conidia trapping and the medium was constantly perfused into the medium infusion channel. This compartmentalization eliminated cross-contamination due to cell-cell interaction and hypha fusion, and subsequently induced growth of the hypha from a narrow channel into the hypha observation channel, allowing an individual hypha to be analyzed independently and accurately.

We evaluated the hyphal extension of *T. reesei* QM9414 in the hyphal growth channels. The channel geometry had great impact on hyphal growth and morphogenesis. To assess the effect of the confinement level, we cultivated *T. reesei* in a device with varying widths (5.0, 7.5, 10.0, 12.5, 15.0, and 17.5 μm). The hyphae growing in wider channels (15 to 20 μm) generated significant branching, which interfered with the visualization of the leading hyphae (Fig. 4B). In contrast, although no branching was observed in hyphae growing in the 5 μm channels, the hyphal morphology (particularly cell borders) was obscured as the hyphae occupied the entire channel width. The length of individual hyphae growing in each channel of QM9414 on the microdevice as a function of time showed that the rate of hyphal elongation across different channel widths was similar (Fig. 4C).

### Organization and movement of the nucleus

The microdevices we constructed could control the growth of a single hypha and allowed for the focused observation under the microscope for long periods. To validate the usefulness of our system to explore intracellular processes, we observed nuclear organization and migration in real time using *T. reesei* QM9414 with histone H2B-GFP nuclear markers. The hyphae of filamentous fungi are composed of multinucleate compartments surrounded by septa, and the proper movement and arrangement of nuclei are crucial for the growth and development of the fungus (26, 27). To investigate how the distribution and organization of nuclei were maintained, we examined the pattern of nuclear migration in a growing hypha. Nuclei moved continuously towards the apical tip of the hypha as it elongated but formed a zone of nuclear exclusion within a certain distance from the apical tip where no nuclei were present (Fig. 5A). The movement of the hyphal tip and of the leading nucleus (closest to the hyphal tip) were linear throughout the period. This suggests that the leading nucleus moves at a similar rate as the hyphal tip (Fig. 5A).

**Figure 5.**
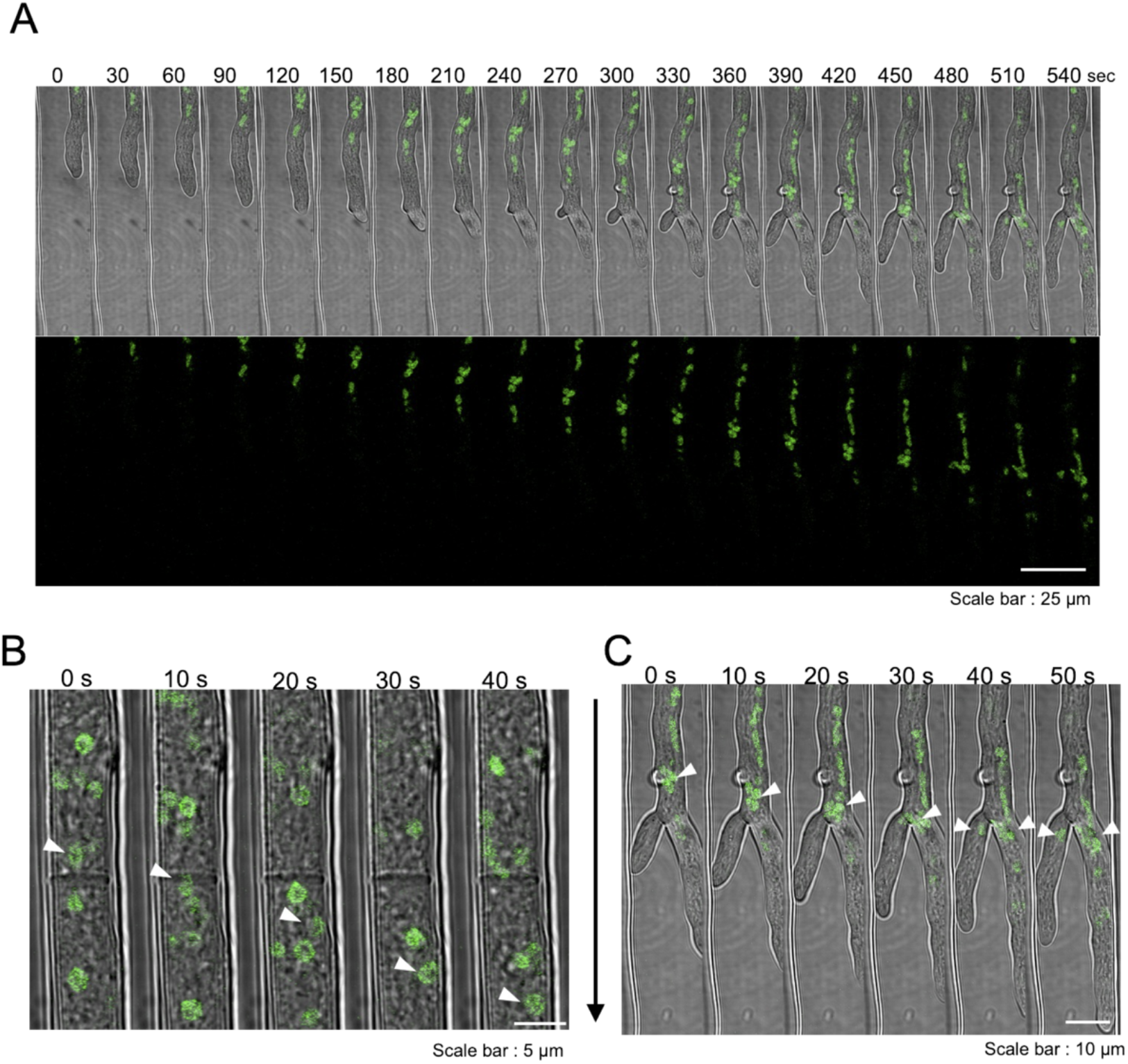
Nuclear dynamics observed with the aid of microdevices. (A) Time-lapse imaging of long-range nuclear migration with combined with bright-field and fluorescence microscopy over 9 min at 30 second intervals. The extension of the hyphal tip, the migration of the leading nucleus, and the distance of the nuclear exclusion zone were observed. Scale bar: 25 μm. The images are from Supplementary Movie S1. (B) Nuclear dynamics at the septum. The images are from Supplementary Movie S2. Scale bar: 10 μm. (C) Nuclear dynamics in hyphal branches. The images are from Supplementary Movie S1. Scale bar: 10 μm. Arrows indicate the direction of hyphal extension.

For further investigation of the local nuclear dynamics, the septum of the young hypha and the bifurcation of the hypha were observed at a higher magnification and in a shorter time. The nucleus was found to pass through the septal pores, with some deformation when it reached the septum (Fig. 5B). When the cluster reached a branch of the hypha, some of the nuclei separated and moved to a new hypha (Fig. 5C).

Furthermore, we also observed characteristic movements of nuclei and vacuoles. Normally, the nucleus moves regularly towards the tip of the hypha by bulk flow in the cytoplasm. However, the fused and enlarged vacuoles pushed the nuclei towards the outer wall of the hypha, delaying their passage through the septa (Fig. 6A). The number of nuclei passing through a particular septum per unit time, and the diameter and number of vacuoles flowing within a single cell were measured. The volume of vacuoles in a particular cell was calculated as a sphere and the percentage of vacuoles in the hypha was calculated (Fig. 6B). At around 130 min when the percentage of vacuoles occupying the hypha increased rapidly, the rate of the movement of nucleus decreased rapidly. This suggested that vacuoles inhibited the regular migration of nuclei. Active nuclear migration (retrograde or accelerated anterograde) was also observed. In contrast to the progressive nuclei (Fig. 6C, red), subsequent nuclei deformed and elongated longitudinally while rapidly retrograding close to the cell wall (Fig. 6C, white).

**Figure 6.**
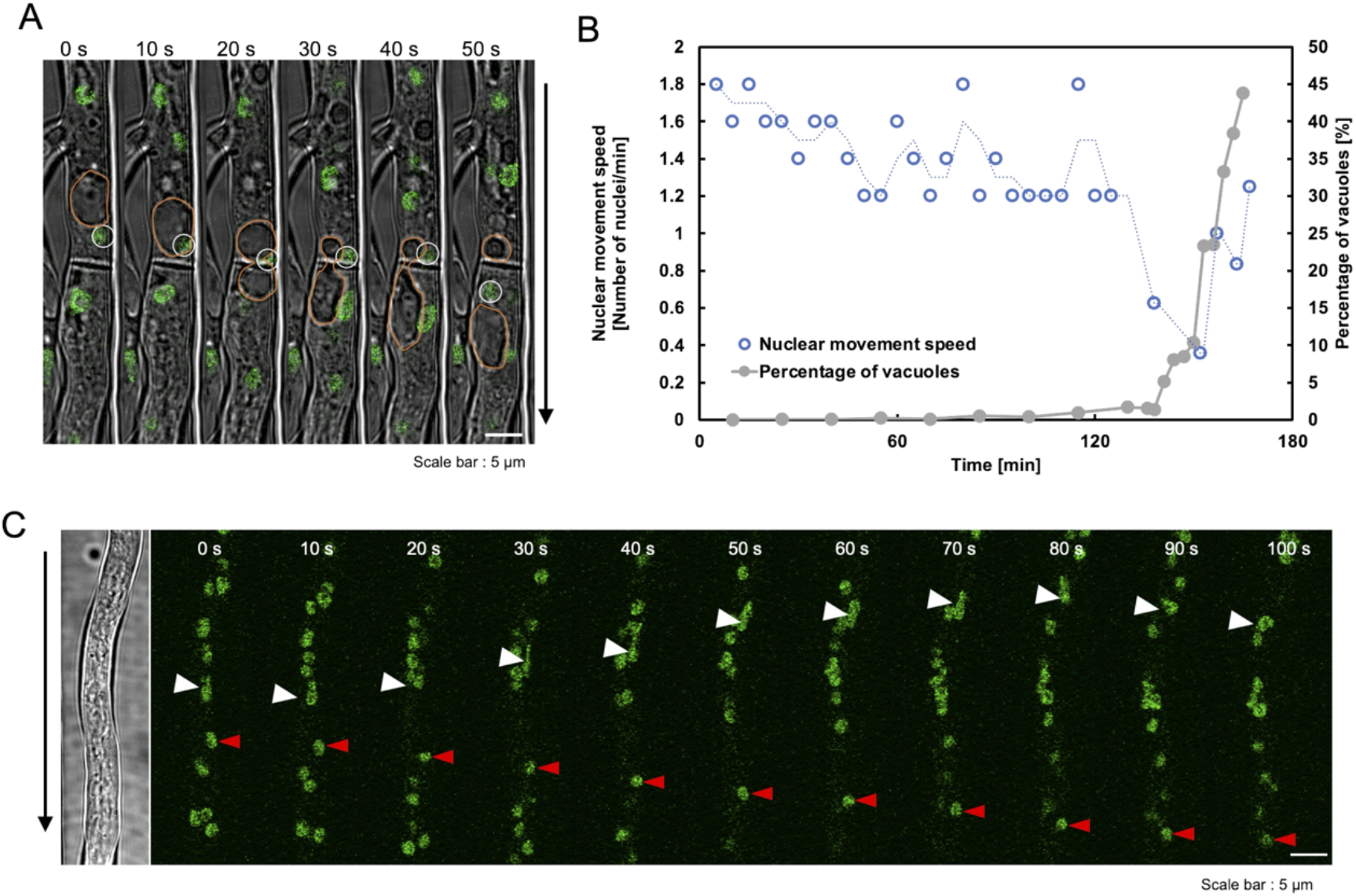
Characteristic movement of nuclei. (A) Nucleus was impeded in its passive movement by a large vacuole; image from Supplementary Movie S3. Scale bar: 5 μm (vacuole: orange, nucleus: white). (B) Vacuoles occupy a proportion of hypha and influence the speed of the movement of the nucleus. Measurements were on the cell from (A). (C) Retrograde nucleus. Supplementary Movie S4. Scale bar: 5 μm (advancing nucleus: red arrows, reverse direction: white arrows).

### Investigation of detection methods for cellulases

To investigate the correlation between hyphal growth and enzyme secretion, we devised a method that detected enzymes inside the device using 4-methylumbelliferyl β-D-cellobioside (4-MUC). 4-MUC is cleaved by cellulase to release 4-methylumbelliferone (4-MU), which emits fluorescence and can be used for the measurement and screening of cellulase activity (Fig. 7A). 4-MU is a coumarin-based fluorescent substance, which is very hydrophobic and is hardly soluble in water. Therefore, it is necessary to use methanol or DMSO to measure the fluorescence intensity. Accordingly, basic mediums containing 4-MUC with or without methanol were prepared and cellulase preparations were added to them. As the degradation product 4-MU is insoluble in water, the fluorescence intensity of the solution with methanol increased, whereas the fluorescence intensity of the solution without methanol did not increase. When the washed mycelia were added to each reaction solution and were observed in preparations, we found that the whole cytoplasm of the fungus fluoresced in the reaction solution without methanol (Fig. S2). No fluorescence was observed in the vacuole or the dead cells (Fig. 7B).

**Figure 7.**
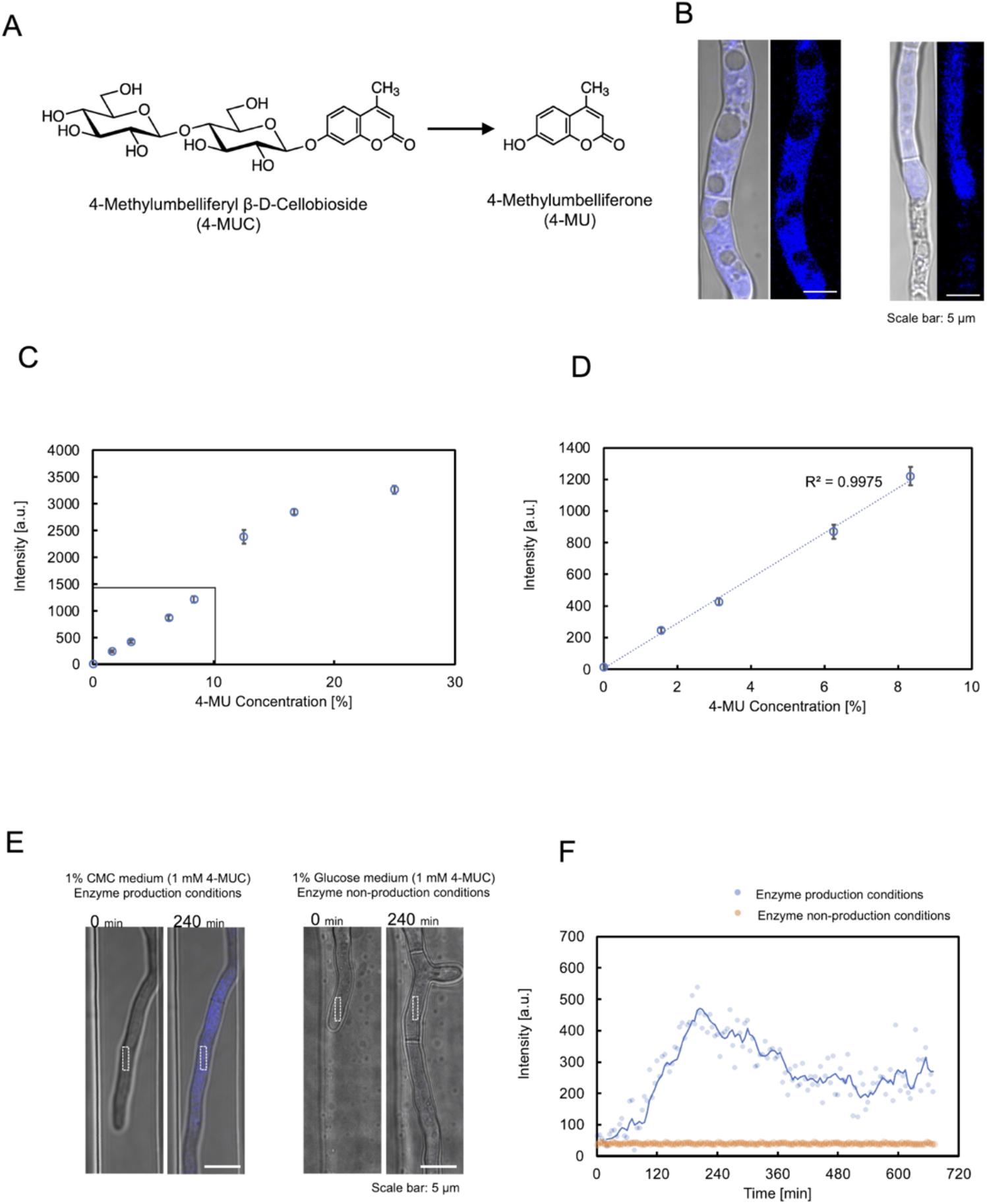
Verification of cellulase detection using 4-MUC. (A) 4-MUC liberates 4-MU by cellulase. (B) 4-MUC and hypha in the enzyme reaction solution. (C) Calibration curve of 4-MU and hyphal fluorescence intensity. (D) Calibration curve of 4-MU and hyphal fluorescence intensity at low concentration. (R^2^ = 0.99) (E) Difference in hyphal fluorescence intensity between enzyme-producing (left) and enzyme-nonproducing (right) conditions (white box: fluorescence measurement site) (F) Time course of fluorescence intensity (E).

We prepared a solution by adding 4-MU to the basic medium and filtering out the remaining crystals that had not dissolved. When the fungus was added to this solution, fluorescence was observed in the cytoplasm. Since the absolute value of the actual concentration of the solution was unknown, we used this solution as a saturated 4-MU solution (100%) and constructed a calibration curve. At low concentrations, there was a linear relationship between 4-MU concentration and fluorescence intensity (R^2^=0.99) (Fig. 7D). The intensity of cytoplasmic fluorescence decreased when the fungus was washed again with basic medium after being exposed to a saturated 4-MU solution. When the fungus was incubated in 1% glucose medium with 1 mM 4-MUC (non-cellulase producing condition), the cytoplasmic fluorescence intensity did not increase at all, whereas when the fungus was incubated in 1% CMC medium with 4-MUC (cellulase producing condition), the fluorescence intensity of the hypha increased over time (Fig. 7E, F). This observation demonstrated that the detection of cellulase by 4-MUC in the micro devise was possible. The growth, morphology and enzyme production in the culture with 4-MUC are shown in Fig. S3.

### Tracking of cellulase dynamics inside and outside the hypha

Finally, we attempted to monitor enzyme production and hyphal behavior. To study the relationship between the dynamics of enzyme production and hyphal morphology, it was necessary to increase the sensitivity of detection of morphological changes, expression of genes encoding enzymes, and of secreted enzymes. We have shown that our device can unravel morphological variations. However, we also encountered a few limitations. *T. reesei* shows the best performance for the enzyme production when solid cellulose is used as a carbon source, whereas only soluble sugars can be used as culture substrates because of clogging in devices with small channels. Therefore, we used the PC-3-7 strain, which has acquired high cellulase inductivity against not only microcrystalline cellulose and sophorose but also cellobiose (6). Furthermore, by fusing GFP to CBHI, the most abundant cellulase produced, we monitored the cellulase expression behavior.

Experiments were carried out steadily for about 48 h. To ensure that each hypha produced the enzyme at the same time, the first incubation period was carried out in a non-enzyme-producing medium (1% glucose, 1 mM 4-MUC). When the hyphal length reached about 150 μm, the medium was switched to an enzyme-producing medium (1% cellobiose, 1 mM 4-MUC). Fluorescence intensity and hyphal growth were measured every 5 minutes. The fluorescence intensity of 4-MU did not match in each channel. This indicated that each channel was compartmentalized, and that the extracellular enzyme profile could be separately measured (Fig. 8A, B, C). The moving average of 4-MU and GFP fluorescence intensities of the same hypha showed a strong correlation (R^2^ = 0.7∼0.9). The hypha continued to grow with some fluctuations and no death or lysis was observed during the observation period (Fig. 8, left).

**Figure 8.**
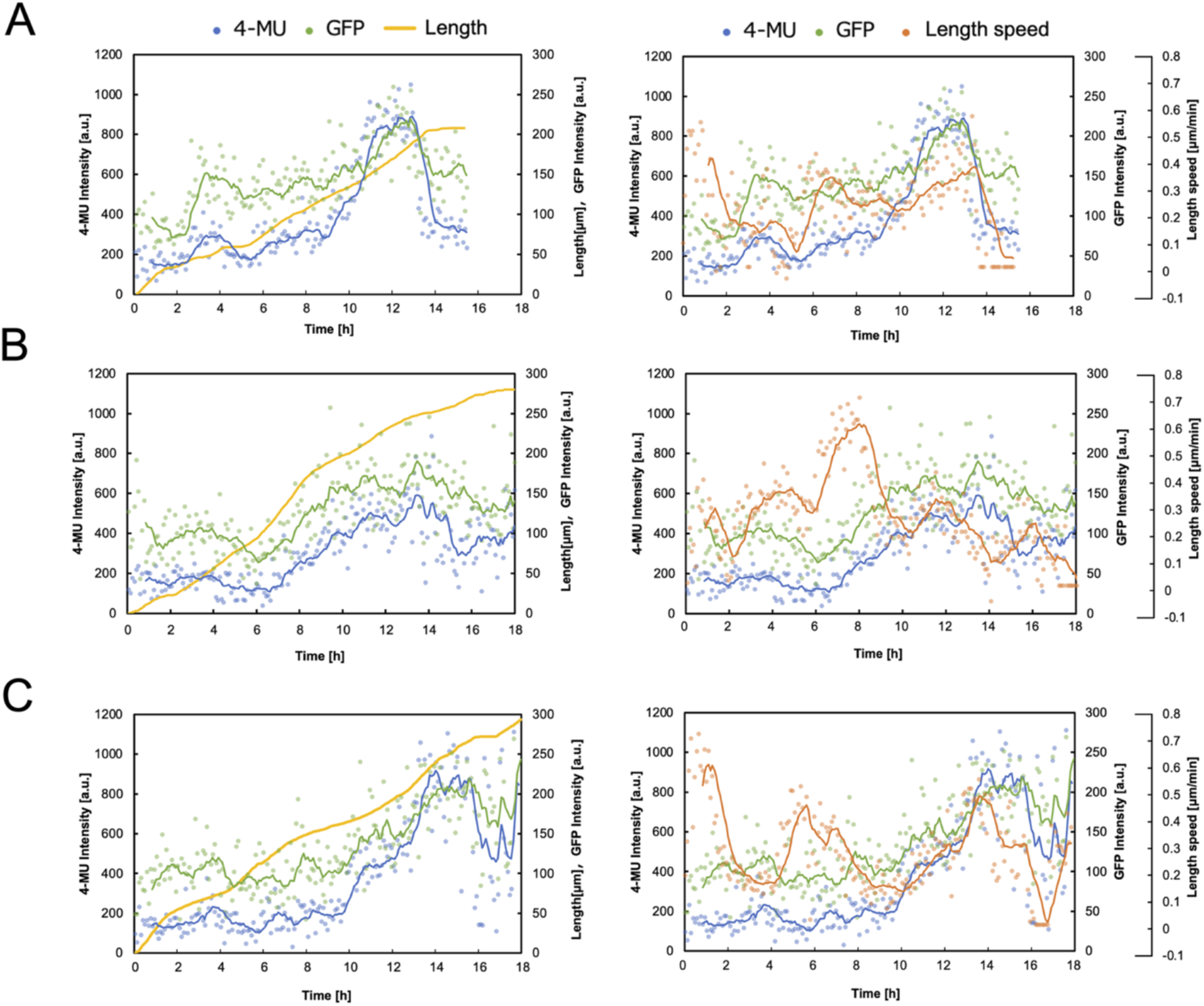
Correlation between the fluorescence intensity of 4-MU and GFP in the hypha and hyphal elongation. Hypha elongation length versus fluorescence intensity is shown on the left and hypha elongation speed is shown on the right. As the fluorescence intensity varied greatly due to the movement of the vacuoles, it is shown as a moving average. (A) Channel width 15 μm. (B) Channel width 10 μm. (C) Channel width 7.5 μm

The enzyme profile showed two characteristic peaks: the first was a low peak that started at about 1 hour after medium exchange and decreased at about 6 hours. In the second peak, the fluorescence intensity increased immediately after the convergence of the first peak and showed a large peak. The molecular diffusion profile of the device (Fig. 3B) showed that the components of the enzyme-producing medium reach the hypha about one hour after the medium change and that a steady state was reached before two hours elapsed. Between 1 and 2 hours the hyphal growth rate uniformly decreased, followed by the first peak in fluorescence intensity. The fluorescence intensity of 4-MU and GFP increased with increasing hyphal elongation rate and decreased with decreasing growth rate or stopping of the hyphal growth. However, the correlation coefficients were low because the hyphal elongation rate did not correspond to the amount of variation in enzyme production.

## DISCUSSION

In this study, we presented a new microfluidic device for the morphological observation and quantification of enzyme production by the filamentous fungus *T. reesei*. Our microfluidic device provides many technical advantages for studying the growth and dynamics of hypha. First, by trapping a single conidium of similar size, each hypha could be divided into isolated compartments, allowing independent and accurate measurements (Fig. 2). In addition, the medium and other components were continuously supplied to the hypha by diffusion (Fig. 3A). Molecular diffusion provides sufficient nutrients for the growing hyphae and the method of medium exchange does not disturb the position of the hyphae, and the method avoids shear stress, which could affect growth studies. Furthermore, by trapping only swollen conidia and expelling conidia that do not germinate or have delayed germination, it was possible to significantly improve the germination rate and to align the germination time (Fig. 4B). The layout of the microfluidic device, which combines these features, allows a stable and robust long-term operation for semi-automatic simultaneous monitoring of different hyphae using multi-point time-lapse.

Previous studies on hyphal dynamics have mainly been carried out on hyphae in agar media, which makes it difficult to observe the hyphae intensively over long periods (34, 35). With this new system, it is now possible to study in detail and in real time, the variability and heterogeneity between cells during hyphal growth, the distribution and dynamics of intracellular organelles, the dynamics of long-term gene expression at different developmental stages in specific locations, and the response of cells to well-defined environmental signals and chemical stimuli.

To validate the usefulness of our system for probing cellular processes, we used *T. reesei* strain QM9414 possessing a histone H2B-GFP nuclear marker that enables the observation of the nuclear organization and movement in real time. The nuclei of *T. reesei* migrated continuously with the elongation of the hypha, forming a nuclear exclusion zone at the tip (Fig. 5A). Hence, the general forward nuclear movement in *T. reesei* is expected to be controlled mainly by the bulk flow of cytoplasm, as observed in previous studies in *N. crassa* (34, 35). In addition, we also observed a characteristic vacuolar behavior. The proportion of the total cell volume occupied by vacuoles varies greatly between fungi and between cells of a particular fungus. Many filamentous fungi have a large number of small vacuoles dispersed in the cytoplasm, but it is also known that some species have much larger vacuoles behind the apical region (36, 37). If *T. reesei* is regulated in the same way as *N. crassa*, the nucleus should move regularly towards the tip of the hypha as shown in Fig. 5. In *T. reesei*, however, the enlarged vacuoles pushed the nuclei towards the outer wall of the hypha and delayed their passage through the septa (Fig. 6A, B). Although there are few reports on the behavior of vacuoles and nuclei in *T. reesei*, and the principle is unknown, the fact that this device allowed us to observe the behavior of the organelles in detail over a long period, from the oldest cells at the base of the hypha to the newest cells at the tip, is a great advance and will greatly facilitate future research on filamentous fungi.

The purpose of this study was to investigate the relationship between the growth of the target hypha and the enzyme, and the method of enzyme detection found in this study depends on a subtle difference in polarity between water and hypha.

4-MU is almost insoluble in water, and it is common to dissolve the substrate in a highly polar solution such as methanol or DMSO for the enzymatic reaction (38). 4-MU has been used in many screening methods and enzyme activity assays (39, 40), but there have been no reports of 4-MU being localized in the cytoplasm. Therefore, when 4-MU is released by the enzymatic reaction, it is localized in the solution which is more polar than the cytoplasm. In our experiments, however, the 4-MU dissolved in a very small amount of water that was concentrated in the cells, which are more polar than water, and that increased the fluorescence intensity of the cytoplasm. When the concentration of methanol in the solution was high, no fluorescence was detected in the hypha, but the solution itself fluoresced. When 4-MU is added to the hypha, the cytoplasmic fluorescence immediately increases. The topological polar surface area value (calculated by Molinspiration Cheminformatics, Slovensky Grob, Slovak Republic) was 208.74 for 4-MUC and 50.44 for 4-MU, indicating that 4-MU has higher membrane permeability. Therefore, it is highly unlikely that 4-MUC is taken into the hypha by the transporter and converted to 4-MU by the intracellular enzyme, and the localization of 4-MU in the cytoplasm is due to membrane permeabilization. In addition, when dead cells with remaining cell walls were stained with Calcofluor White, only the cell walls were stained, whereas when 4-MU alone was added to the same cells, the cell walls were not stained, indicating that 4-MU was not attached to the cell walls.

The hydroxy group of 4-MU facilitates the chemical synthesis of various fluorescently labeled substances. Typical examples are 4-methylumbelliferyl phosphate (4-MUP), 4-methylumbelliferyl-β-D-glucopyranoside (4-MUG) and 4-methylumbelliferyl acetate (MU-Ac). These are used for various purposes. Importantly, 4-MU localizes to the cytoplasm and can be detected even in trace amounts without the adverse effects of methanol or DMSO on the hyphae. Hence the increased detection sensitivity means that reactions missed by previously used fluorescent substrates can now be detected immediately.

We investigated the relationship between hyphal growth and enzyme expression using the novel device and the method we developed to detect extracellular enzymes by 4-MUC. First, the fluorescence intensity of 4-MU did not match in each channel, indicating that the extracellular enzyme profile of each hypha could be measured separately (Fig. 8). Although it is not possible to observe only one hypha because its morphology depends on the strain, culture medium and surrounding geometry, it was possible to measure the growth of hypha in each compartment by measuring only the hypha that grew from a single conidium. The PC*cbh1gfp* strain does not secrete GFP-CHB1. It is known that unsecreted GFP is not degraded intracellularly and accumulates in the vacuole (34, 41). When we sampled flask cultures and observed them every 12 hours, we found that GFP accumulated in the vacuole after 62 hours, but we could not find any organelle with a specific increase in GFP fluorescence intensity before that time.

The fact that we did not observe any vacuoles, cessation of hyphal growth, or lysis during the 20-hour observation period after switching the culture substrate suggests that the GFP fluorescence intensity indicates the amount of CBHI produced in situ for a short period after the *cbh1* promoter was activated. The set of cellulases of *T. reesei* is coordinately expressed and the proportion of each expression amount does not change significantly with different substrates of induction (42). Assuming that the proportion of cellulase produced by the PC*cbh1gfp* strain remained unchanged, the 10-interval mean trends of 4-MU and GFP fluorescence intensities in the same hypha showed a strong correlation (R^2^ = 0.8∼0.9), indicating that we can reliably observe the amount of CBHI and secreted enzymes in the hypha (Fig. 8).

The growth of the fungus is stopped or slowed down when the culture substrate or culture environment changes. Throughout the observation period, the fluorescence intensity of 4-MU and GFP increased with increasing hyphal growth rate and decreased with decreasing growth rate or stopping of hyphal growth, indicating that exocytosis provides both cell wall components and secreted enzymes in real-time. However, the hyphal growth rate and enzyme production pattern were not significantly different. The rate of hyphal elongation did not coincide with the range of variation in enzyme production, suggesting that the amount of enzyme in the secretory vesicle is not constant, but more studies are needed to corroborate the possibility of inconsistent variation due to hyphal branching.

4-MUC is a reagent for cellulase detection and is coordinately degraded by CBH and BGL. *T. reesei* also secretively produces a set of cellulases (42). Cellulase production could be monitored effectively in this study using the new device. This research offers a novel solution to the challenges in the measurement of fungal growth and morphology, enzyme production, and specific promoter strength, all of which have been difficult to measure simultaneously. The microfluidic device that we have developed can measure these parameters simultaneously and in real time. We have also shown that subtle differences in polarity between cells and extracellular solutions can enable the detection of small amounts of enzymatic reactions (Fig. 9).

**Figure 9.**
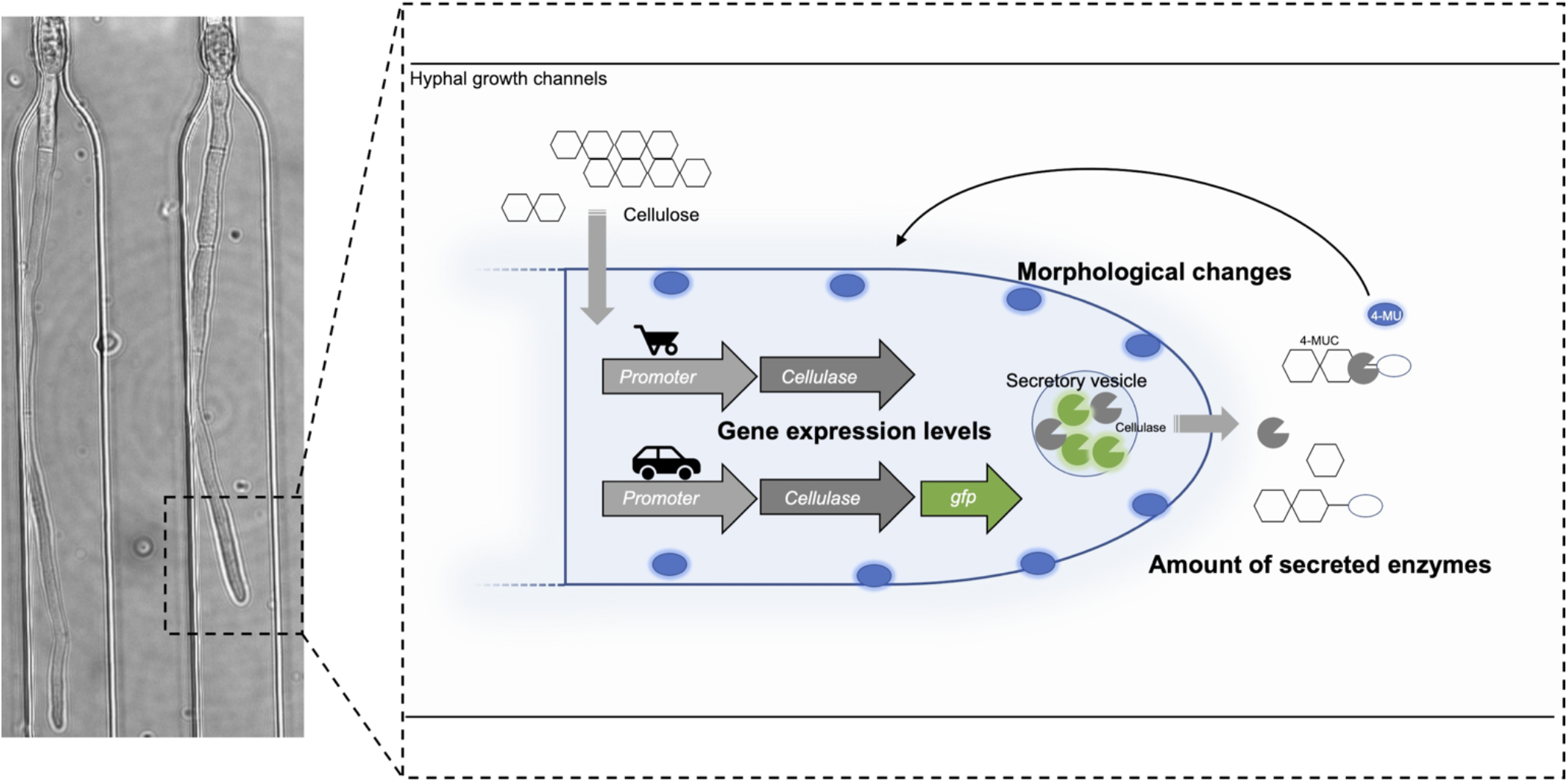
Schematic diagram of growth and morphology, enzyme production and specific promoter strength measurements in *T. reesei*

Further new research areas for the future include improved layout for the microfluidic device, the integration of faster stage movement motors, automated image acquisition microscopy systems, optics and detectors with the higher spatial and temporal resolution, and development of a variety of fluorescent molecular markers. As such, the microfluidic device can also be used to study more complex fungal behavior, such as the branching and fusing of different hyphae within a fungus, by branching and connecting channels in the middle. Furthermore, the integration of multiple loading channels on a single chip allows the comparison of different fungal strains and the study of interactions between fungi with different genotypes. In addition to detection by imaging, techniques such as laser microdissection can be combined to accurately extract sections of hypha from the hypha of interest. In addition, the equipment and strategies presented here can be immediately extended to a variety of other cell types, allowing a wide range of new studies to reveal the characteristics and mechanisms of long-term cell growth.

## MATERIALS AND METHODES

### Fungal strains and growth conditions

*T. reesei* strains QM9414 (ATCC26921) and PC-3-7 (ATCC66579) used in this study were obtained from Kao Co., Ltd. Previously described *pyr4* disruptants QM9414ΔP and PC-3-7ΔKP (28) were used as the host for transformation. Strains were grown on potato dextrose agar (PDA) plates and harvested conidia. Molecules of *T. reesei* strains were inoculated into basic medium (1% appropriate carbon source, 0.14% (NH_4_)_2_SO_4_, 0.2% KH_2_PO_4_, 0.03% CaCl_2_•2H_2_O, 0.03% MgSO_4_•7H_2_O, 0.05% Bacto Yeast Extract, 0.1% Bacto Peptone, 0.1% (w/v) Tween 80, 0.1% (w/v), 50 mM tartrate buffer (pH4.0), and trace element). Trace element comprised 0.006% H_3_BO_3_, 0.026% (NH_4_)6Mo_7_O_24_•4H_2_O, 0.1%FeCl_3_•6H_2_O, 0.4% CuSO_4_•5H_2_O, 0.008% MnCl_2_•4H_2_O, 0.2% ZnCl_2_.

### Biochemical analyses

To obtain the pattern of secreted protein, the culture supernatant was subjected to SDS-PAGE (29). The Precision Plus Dual Color Standard Marker (Bio-Rad) was used as molecular weight standard. Protein concentration was determined by Bradford method using bovine gamma globulin as the standard (30).

### Construction of plasmids and strains

For the GFP-H2B expression, the gene encoding H2B (*h2b*) and its 2 kbp upstream and downstream regions were amplified by PCR with the genomic DNA derived from QM9414 as the template. This fragment was inserted into the HinCII site of the pUC118 vector by the Gibson Assembly system (New England Biolabs) to create pU*h2b*. An inverse PCR was then performed for the linearization of pU*h2b* at the start codon of *h2b* and the gene encoding GFP was introduced into just upstream of *h2b*. The resulting plasmid, pU*h2b-gfp*, was subjected to an inverse PCR to open the downstream region of *h2b*, and the selection marker gene *pyr4* was inserted to obtain plasmid pU*h2b-gfp-pyr4*. The GFP-H2B expression cassette was released from pU*h2b-gfp-pyr4* by XbaI and SbfI. This expression cassette was used to transform the QM9414ΔP strain.

For the CBHI-GFP expression, the gene encoding CBHI (*cbh1*) and its 2 kbp upstream and downstream regions were amplified by PCR. This fragment was inserted into the EcoRV site of the pBluescriptII KS(+) vector by the Gibson Assembly system to create pB*cbh1*. An inverse PCR was then performed for the linearization of pB*cbh1* at the C-terminus of *h2b* and the selection marker gene *pyr4* was introduced into the 1.5 kbp downstream region of *cbh1*. The resulting plasmid, pB*cbh1-pyr4*, was subjected to an inverse PCR to open the downstream region of *cbh1*, and the gene encoding GFP was inserted to obtain the plasmid pB*cbh1-gfp-pyr4*. The VBH1-GFP expression cassette was released from pB*cbh1-gfp-pyr4* by ClaI and SpeI. Using this expression cassette, PC-3-7ΔKP strain was transformed and PC*cbh1gfp* strain was obtained. The primers used in this study are shown in Supplemental Table S1.

### Fungal transformation

Transformation of *T. reesei* protoplasts was carried out by the protoplast-PEG method as previously described (31) with a small modification in which Yatalase (Takara bio) was used for cell wall digestion for preparing the protoplasts. Protoplasts were plated on minimal medium without uridine to screen for cells with uridine autotrophy. Candidates of transformants were streaked twice on minimal medium without uridine to obtain stable transformants. Recombination of strains was confirmed by Southern hybridization analysis using an AlkPhos Direct kit (Cytiva).

### Microfluidic device fabrication

The microfluidic devices were fabricated by standard photolithographic techniques (details in Fig. S1) (32, 33). The molds for the PDMS devices were fabricated in two steps using SU-8 3205 and 30025 on silicon wafers. PDMS (SYLGARD 184 Silicone Elastomer Kit) was poured into the SU-8 mold in a container and allowed to cure. Holes were drilled and bonded together with coverslips (24× 60 mm, Matsumani) activated with a plasma cleaner (PDC-32G, Harrick Plasma).

### Operation of the microfluidic device

*T. reesei* conidia were suspended in basic medium containing 1% glucose and left to swell at 28°C for about 5 hours. The conidia obtained by centrifugation were then suspended in basic medium and injected into the microfluidic device. Then, the basic medium containing appropriate carbon source (1%) and 1 mM 4-MUC was constantly perfused into the microfluidic device at a flow rate of 0.5 mL/h using a syringe pump (YSP-301, YMC).

### Image acquisition and analysis

The microfluidic device was mounted on the stage of a Nikon Ti2-C2 confocal microscope equipped with 60× (Apo Lambda S Oil, Nikon) and 100× (Apo TIRF Oil, Nikon) objective lenses. Crosstalk was minimized by the use of line sequential mode. The images were stored and manually analyzed using the Nikon NIS-Elements software, which is compatible with the microscope.

### Characterization of molecular diffusion

The microfluidic device was first filled with basic medium, and then perfused with fluorescent staining reagents Calcofluor White (CFW; Sigma) solution into the medium infusion channel using a syringe pump. Fluorescence images of hyphal growth channels were captured after CFW introduction, and the intensity was quantified.

## ACKNOWLEDGMENTS

We thank Y. Wakamoto for the MEMS technical assistance.

This study is based on results obtained from a project, JPNP19001, commissioned by the New Energy and Industrial Technology Development Organization (NEDO).

A.I., Y.S., and W.O. helped in conceptualization; A.I. performed the research; W.O. contributed new reagents/analytic tools; W.O. contributed to funding acquisition; Y.S., and W.O. supervised; and A.I., Y.S., and W.O. wrote the paper. All authors discussed the results and commented on the manuscript.

## Supplemental Material

Table S1

Primers used in this study. Genomic DNA from QM9414 was used as a template to amplify the target gene. pUC118 vector was used for GFP-H2B expression and pU*h2b-gfp-pyr4* was created. the target gene was amplified in the same way for CBHI-GFP and pBluescriptII KS(+) Inserted into a vector to create pB*cbh1-gfp-pyr4*.

Fig S1

Microfluidic device fabrication procedures.

Fig S2

Differences in 4-MU fluorescence with and without methanol. When the fluorescent substance 4-MU was dissolved in 5% methanol, the solution fluoresced dimly, but the hypha did not. When 4-MU was added to the medium, the solution did not fluoresce, but the hypha fluoresced strongly.

Fig S3

(A to D) The growth, morphology and enzyme production in the culture with 4-MUC. Comparisons were made between 1% glucose medium with 1 mM 4-MUC added to it and 1% cellobiose medium with 1 mM 4-MUC added to it. (A) Dry cell weight transition. (B) Extracellular protein concentration at 48 hours of incubation. (C) SDS-PAGE of culture supernatant from (B). (D) Growth in 4-MUC-supplemented culture medium. Taken at 24 h of incubation. Scale bar: 100 μm.

Movie S1

Time-lapse imaging of long-range nuclear migration with combined with bright-field and fluorescence microscopy at intervals of 30 second intervals. The extension of the hyphal tip, the migration of the leading nucleus, and the distance of the nuclear exclusion zone were observed. Scale bar: 25 μm.

Movie S2

Time-lapse imaging of long-range nuclear migration with combined with bright-field and fluorescence microscopy at intervals of 10 second intervals. Nuclear dynamics at the septum. Scale bar: 10 μm.

Movie S3

Time-lapse imaging of long-range nuclear migration with combined with bright-field and fluorescence microscopy at intervals of 10 second intervals. Nucleus was impeded in its passive movement by a large vacuole. Scale bar: 5 μm

Movie S4

Time-lapse imaging of long-range nuclear migration with combined with bright-field and fluorescence microscopy at intervals of 10 second intervals. Retrograde nucleus. Scale bar: 5 μm (reverse direction: white circle).

